# Exposure to a Healthy Gut Microbiome Protects Against Reproductive and Metabolic Dysregulation in a PCOS Mouse Model

**DOI:** 10.1101/472688

**Authors:** Pedro J. Torres, Bryan S. Ho, Pablo Arroyo, Lillian Sau, Annie Chen, Scott T. Kelley, Varykina G. Thackray

## Abstract

Polycystic ovary syndrome (PCOS) is a common endocrine disorder affecting approximately 10% of reproductive-aged women worldwide. Diagnosis requires two of the following: hyperandrogenism, oligo/anovulation and polycystic ovaries. In addition to reproductive dysfunction, many women with PCOS display metabolic abnormalities associated with hyperandrogenism. Recent studies have reported that the gut microbiome is altered in women with PCOS and rodent models of the disorder. However, it is unknown whether the gut microbiome plays a causal role in the development and pathology of PCOS. Given its potential role, we hypothesized that exposure to a healthy gut microbiome would protect against development of PCOS. A co-housing study was performed using a letrozole-induced PCOS mouse model that recapitulates many reproductive and metabolic characteristics of PCOS. Since mice are coprophagic, co-housing results in repeated, non-invasive inoculation of gut microbes in co-housed mice via the fecal-oral route. In contrast to letrozole-treated mice housed together, letrozole-treated mice co-housed with placebo mice showed significant improvement in both reproductive and metabolic PCOS phenotypes. Using 16S rRNA gene sequencing, we observed that the gut microbial composition of letrozole-treated mice co-housed with placebo mice differed from letrozole mice housed together. In addition, our analyses identified several bacterial taxa including *Coprobacillus, Dorea* and *Adlercreutzia* associated with the improved PCOS phenotype in letrozole-treated mice co-housed with placebo mice. These results indicate that disruption of the gut microbiome may play a causal role in PCOS and that manipulation of the gut microbiome may be a potential treatment option for PCOS.

**Significance:** PCOS is a common cause of female infertility and ~80% of women with PCOS have metabolic dysregulation that predisposes them to type 2 diabetes and cardiovascular disease. Since treatment options for the metabolic symptoms of PCOS are limited, there is a need to develop novel therapeutic options. The gut microbiome has emerged as an important player in human health and has been shown to play a causal role in obesity. In this study, we found that exposure to a healthy gut microbiome through co-housing protected against the development of reproductive and metabolic dysregulation in a PCOS mouse model. These results suggest that manipulation of the gut microbiome may be a potential treatment option for women with PCOS.

## Introduction

Polycystic ovary syndrome (PCOS) is a common endocrine disorder affecting approximately 10% of women worldwide (1). Diagnosis of PCOS, using the Rotterdam Consensus criteria (2003), requires two of the following: hyperandrogenism, oligo-or amenorrhea and polycystic ovaries. PCOS is the leading cause of anovulatory infertility in women, and women with PCOS also have an increased likelihood of miscarriage and pregnancy complications. Though it is often perceived as a reproductive disorder, PCOS is also a metabolic disorder. Women with PCOS have an increased risk of developing obesity, type 2 diabetes, hypertension, and non-alcoholic fatty liver disease (2–4). PCOS-related metabolic dysfunction is associated with hyperandrogenism and occurs irrespective of body mass index (5, 6). While studies indicate that androgen excess is an important contributor to metabolic dysregulation in women with PCOS, the mechanisms that lead to obesity and insulin resistance in PCOS are not well understood. Although genetic and environmental factors undoubtedly influence the development and pathology of PCOS (7–10), it is worth exploring whether gut microbes contribute to this disorder.

Studies over the past decade have shown that the gastrointestinal tract harbors a complex microbial ecosystem (the gut microbiome) that is important for human health and disease (11, 12). Gut microbes offer many benefits to the host including protection against pathogens as well as regulation of host immunity and the integrity of the intestinal barrier (13–15). The gut microbiome is also involved in the production of short-chain fatty acids via fermentation of dietary fibers, production of essential vitamins such as folic acid and B12 and modification of bile acids, neurotransmitters and hormones (16, 17). Studies have also shown that changes in the gut microbiome are associated with metabolic disorders such as obesity and type 2 diabetes (18, 19). Moreover, studies have reported that transplantation of stool from obese donors into germ-free mice results in an obese phenotype (20), suggesting that the gut microbiome may play a causative role in metabolic dysregulation. These transplantation studies were complemented with co-housing studies that took advantage of the fact that, since mice are coprophagic, co-housing provides a means for repeated, non-invasive microbial inoculation. Co-housing germ-free mice transplanted with stool from obese donors with germ-free mice transplanted with stool from lean donors was shown to protect the mice transplanted with obese donor stool from developing obesity (20–22). Altogether, these studies suggest that modulation of the gut microbiome may be a potential treatment option for metabolic disorders.

With regards to PCOS, several recent studies reported that changes in the gut microbiome are associated with PCOS (23–26). These studies detected lower alpha diversity and differences in the relative abundances of specific Bacteroidetes and Firmicutes in women with PCOS compared to control individuals (23–25). In particular, decreases in the relative abundance of *Bacteroides* and genera from the Ruminococcaceae and S24-7 families were observed in several studies. In addition, changes in the gut microbiome correlated with hyperandrogenism (25), suggesting that testosterone may influence the composition of the gut microbiome in females. In addition to studies in humans, several studies reported a significant association between the gut microbiome and PCOS in rodent models (27, 28). Since the rodent models are diet independent, these studies suggest that the mechanisms that result in an altered gut microbiome in PCOS are distinct from diet-induced effects on the gut microbiome observed in obesity models.

We previously developed a PCOS mouse model that uses treatment with letrozole, a non-steroidal aromatase inhibitor, to increase testosterone levels and decrease estrogen levels by inhibiting the conversion of testosterone to estrogen (29). Letrozole treatment of pubertal female mice results in the recapitulation of many reproductive hallmarks of PCOS including hyperandrogenism, acyclicity, polycystic ovaries, and elevated luteinizing hormone (LH) levels. This model also exhibits metabolic dysregulation similar to the phenotype in women with PCOS including weight gain, abdominal adiposity, increased fasting blood glucose (FBG) and insulin levels, impaired glucose stimulated-insulin secretion, insulin resistance, and dyslipidemia (30). A caveat is that low estrogen levels in this model do not reflect estrogen levels in women with PCOS. Similar to women with PCOS, 16S rRNA gene sequencing showed that letrozole treatment was associated with lower microbial richness of the gut microbiome, a shift in the overall gut microbial composition, and changes in specific Bacteroidetes and Firmicutes. Since a recent study examining the effects of non-antibiotic drugs on the gut microbiome found that letrozole did not alter growth of ~40 representative gut bacteria (31), this suggests that the differences in gut microbial composition found in the PCOS mouse model are not due to a direct effect of letrozole. Overall, these studies indicate that the gut microbiome of individuals with PCOS differs significantly from healthy individuals and suggest that a microbial imbalance or “dysbiosis” in the gut may contribute to the pathology of PCOS.

To begin to address whether the gut microbiome contributes to PCOS pathophysiology and if manipulation of the gut microbiome can be used to treat PCOS, we used a co-housing paradigm to determine whether exposure to a healthy gut microbiome protected against development of PCOS metabolic and reproductive phenotypes. Since mice are coprophagic, gut microbes can be readily transferred from one mouse to another through the fecal-oral route. In this study, pubertal female mice were treated with placebo or letrozole and co-housed two per cage for 5 weeks. The groups consisted of co-housed placebo mice, co-housed letrozole mice or a placebo mouse cohoused with a letrozole mouse. Overall, co-housing letrozole with placebo mice resulted in substantial improvement in both PCOS metabolic and reproductive phenotypes compared to letrozole mice co-housed together. Furthermore, 16S rRNA gene sequence analysis demonstrated that co-housing letrozole with placebo mice resulted in changes in the beta diversity of the gut microbiome and highlighted specific bacterial genera that may be candidates for pre- or probiotic therapies. Our findings support the idea that there may be a causal link between the gut microbiome and PCOS and that restoration/manipulation of the gut microbiome may be a potential treatment option for PCOS.

## Results and Discussion

### Co-housing letrozole mice with placebo mice resulted in less weight gain and abdominal adiposity

To investigate whether exposure to a healthy gut microbiome can protect against the development of a PCOS metabolic and reproductive phenotype, we performed a co-housing study. Female mice were implanted with a placebo or letrozole pellet at 4 weeks of age (n=8/group). The mice were divided into 3 groups and housed two mice per cage resulting in 4 groups: co-housed placebo mice (P/P), co-housed letrozole mice (L/L) and co-housed placebo with letrozole mice (P/L and L/P, respectively) (Fig. 1A). As shown in Fig. 1B, weight was measured each week during the 5 weeks of treatment. Similarly to previously published studies (27, 30), 2 weeks of letrozole treatment resulted in increased weight compared to placebo treatment that was maintained for the rest of the study (Fig. 1B). Five weeks of letrozole treatment also increased abdominal adiposity compared to placebo treatment (Fig. 1C). Interestingly, P/L mice had similar weight gain and abdominal adiposity compared to P/P mice. In contrast, L/P mice gained less weight and exhibited a trend towards less abdominal adiposity compared to L/L mice (Fig. 1B-C). It is noteworthy that our previous study showed that letrozole treatment induced weight gain and abdominal adiposity in female mice without a change in food intake or total energy expenditure (30). Since studies have indicated that obesity may be influenced by an increased capacity of the gut microbiome to harvest energy from dietary fiber (21), it would be informative to test whether this occurs in the letrozole-treated gut microbiome and whether exposure to a healthy gut microbiome ameliorated this effect.

**Fig. 1.**
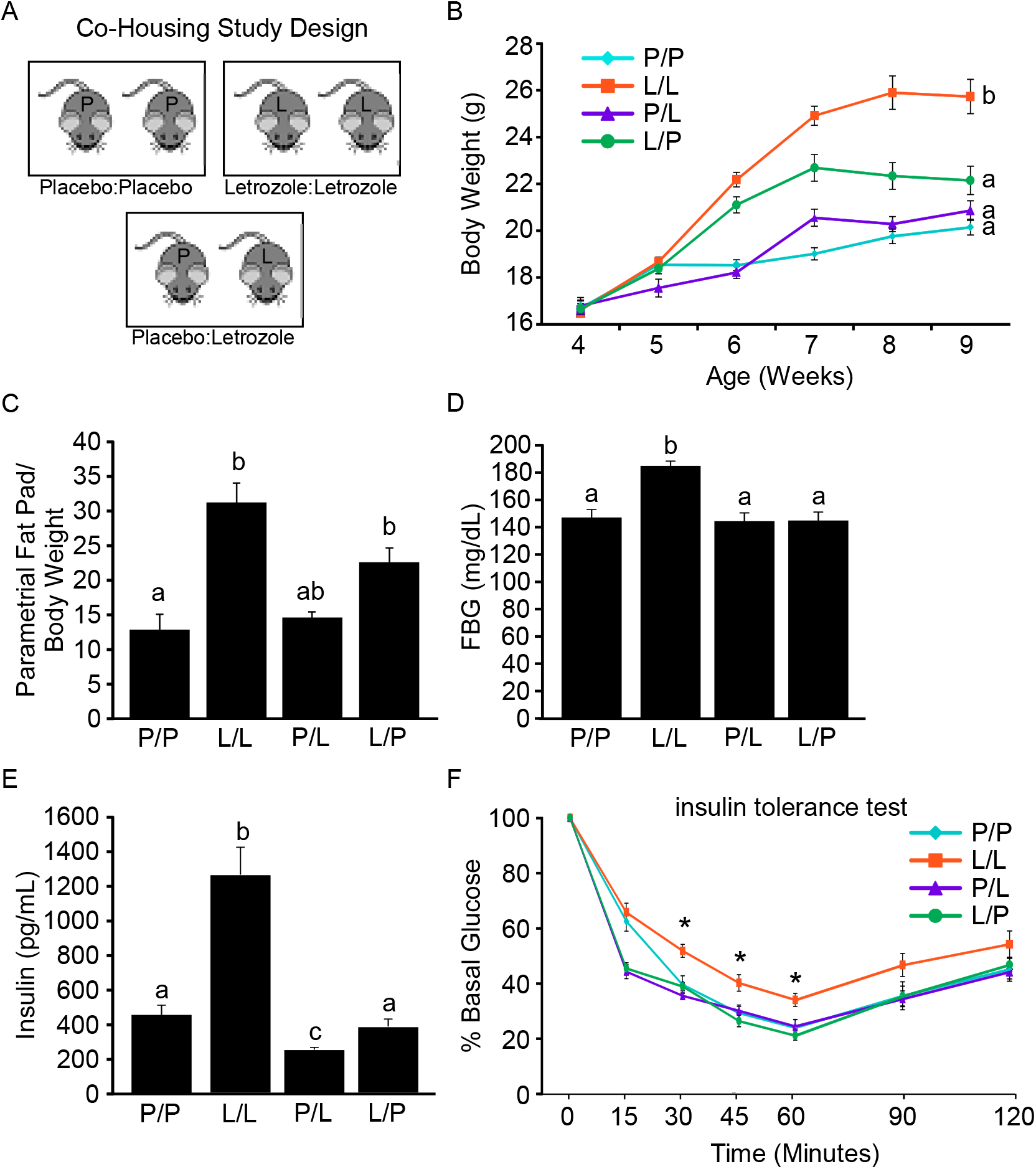
Co-housing letrozole mice with placebo mice protected against development of the PCOS metabolic phenotype. Design of co-housing study over the course of a 5-week experiment included 3 groups of 4-week-old female mice housed 2 per cage: co-housed placebo mice (P/P), co-housed letrozole mice (L/L), and co-housed placebo with letrozole mice (P/L and L/P, respectively) (A). Letrozole treatment (L/L) resulted in metabolic dysfunction compared to placebo (P/P) including increased weight, abdominal adiposity, fasting blood glucose (FBG) and insulin levels and insulin resistance (B-F). Compared to L/L mice, L/P mice showed a decrease in body weight, a decrease in abdominal adiposity, a decrease in FBG and insulin levels, and restored insulin sensitivity (B-F).

### Co-housing letrozole mice with placebo mice resulted in reduced fasting blood glucose and insulin levels and insulin sensitivity

As reported in previous studies (27, 29, 30), five weeks of letrozole treatment resulted in increased FBG and insulin levels and insulin resistance (Fig. 1D-F). P/L mice had similar serum glucose and insulin levels and insulin sensitivity to that of the P/P mice while L/P mice had reduced FBG and insulin levels as well as less insulin resistance compared to L/L mice (Fig. 1D-F). Interestingly, the protective effect of co-housing letrozole mice with placebo mice on weight gain only manifested after several weeks of treatment. While our results demonstrated that cohousing letrozole mice with placebo mice resulted in protection from changes in FBG and insulin levels and insulin resistance by the end of the study, future studies will be required to ascertain how much time co-housing takes to exert a protective effect on these metabolic factors and what triggers these changes.

### Co-housing letrozole mice with placebo mice resulted in estrous cyclicity

In addition to characterizing the effect of co-housing on the PCOS metabolic phenotype, we also assessed the effect on the reproductive axis. As previously published (27, 29), letrozole treatment resulted in hallmarks of PCOS including elevated testosterone and LH levels and acyclicity in the L/L mice (Fig. 2A-C). P/L mice did not have changes in testosterone, LH or estrous cyclicity compared to P/P mice (Fig. 2A-C). On the other hand, L/P mice had decreased testosterone and LH levels compared to L/L mice (Fig. 2A-B). In addition, the L/P mice displayed a full estrous cycle or changes in the morphology of vaginal epithelial cells compared to the constant diestrus exhibited by L/L mice (Fig. 2C).

**Fig. 2.**
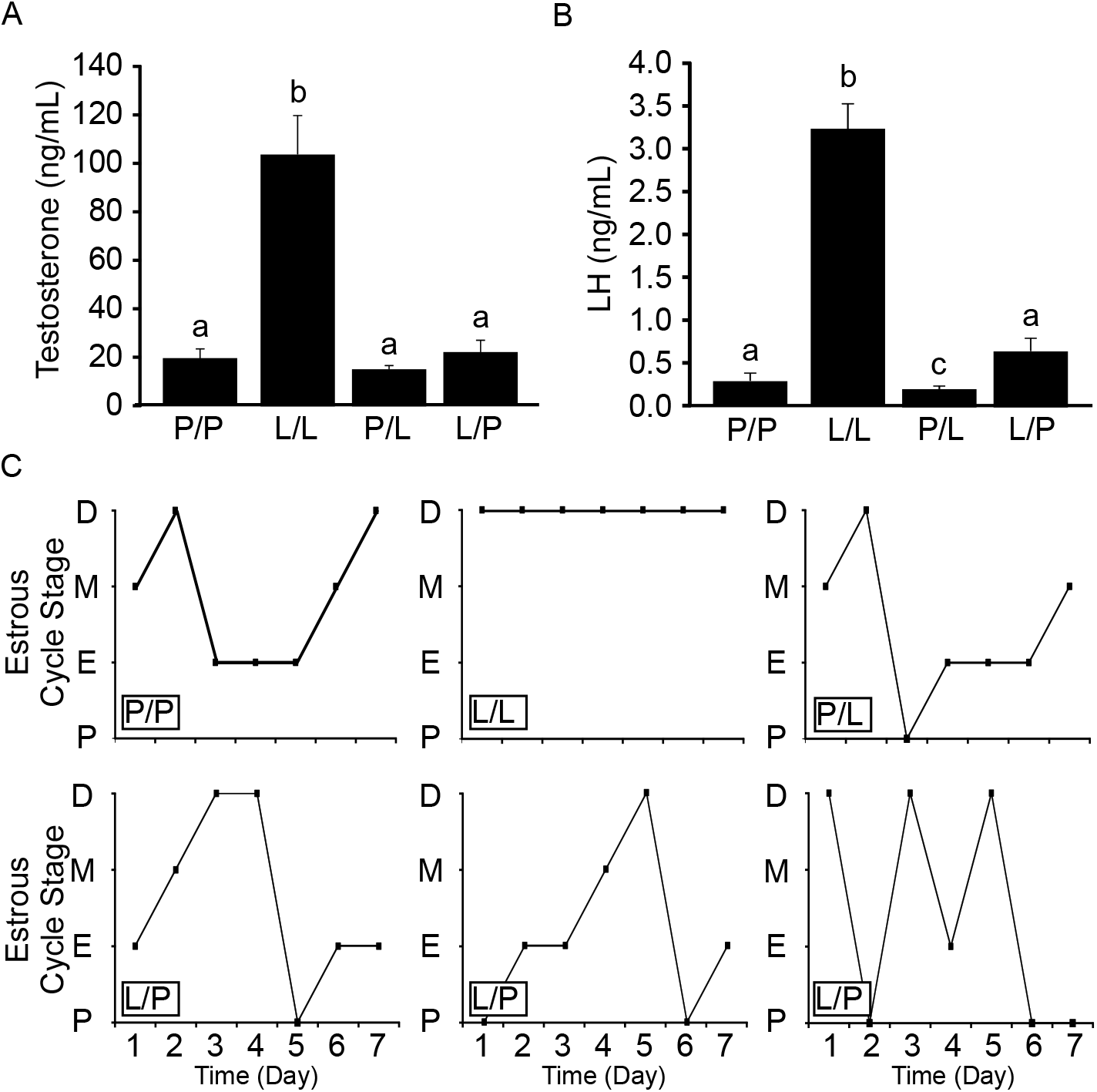
Letrozole mice co-housed with placebo mice did not become hyperandrogenemic or acyclic. The co-housing study included 3 groups of female mice housed 2 per cage: co-housed placebo mice (P/P), co-housed letrozole mice (L/L), and co-housed placebo with letrozole mice (P/L and L/P, respectively). Letrozole treatment (L/L) resulted in increased testosterone and luteinizing hormone (LH) levels compared to placebo (P/P) (A-B). L/P mice displayed a decrease in testosterone and LH (A-B) as well as a restoration of estrous cyclicity compared L/L mice stuck in diestrus (C). Stages of the estrous cycle are indicated as diestrus (D), metestrus (M), estrus (E) and proestrus (P).

### Co-housing letrozole mice with placebo mice protected ovarian function

Consistent with previous reports (27, 29), the ovaries of letrozole-treated mice (L/L) lacked corpora lutea and displayed cystic follicles and hemorrhagic cysts while the ovaries of P/L mice had a similar morphology to that of P/P mice (Fig. 3A). Interestingly, the ovaries of L/P mice lacked cystic follicles and hemorrhagic cysts and contained corpora lutea (Fig. 3A). The presence of estrous cycles and corpora lutea in the L/P mice suggests that the mice were able to ovulate. Similar to previous reports (29), L/L mice showed a significant increase in both ovarian weight and ovarian mRNA expression levels of follicle-stimulating hormone receptor *(Fshr),* cytochrome P450 17A1 *(Cyp17),* and aromatase *(Cyp19)* compared to P/P mice (Fig. 3B-E). The ovarian weight and mRNA expression levels in P/L mice mirrored those of the P/P mice (Fig. 3B-E). Compared to L/L mice, L/P mice showed a significant decrease in ovarian weight and mRNA expression levels of *Cyp17* while *Fshr* and *Cyp19* mRNA expression levels were comparable to L/L mice (Fig. 3B-E). Since *Cyp17* gene expression is induced by both androgen and insulin (32, 33), it is unsurprising that *Cyp17* levels were normalized in L/P mice that had reduced circulating levels of testosterone and insulin. On the other hand, it is not clear why *Fshr* mRNA levels were increased in both L/L and L/P mice. With regards to aromatase expression, it is notable that letrozole treatment resulted in an increase in mRNA levels that was not altered by co-housing letrozole with placebo mice. This data indicates that suppression of the aromatase enzyme with letrozole treatment in both L/L and L/P mice resulted in a compensatory increase in *Cyp19* mRNA and provides indirect evidence that the effect of co-housing did not occur because of decreased letrozole activity in the L/P mice.

**Fig. 3.**
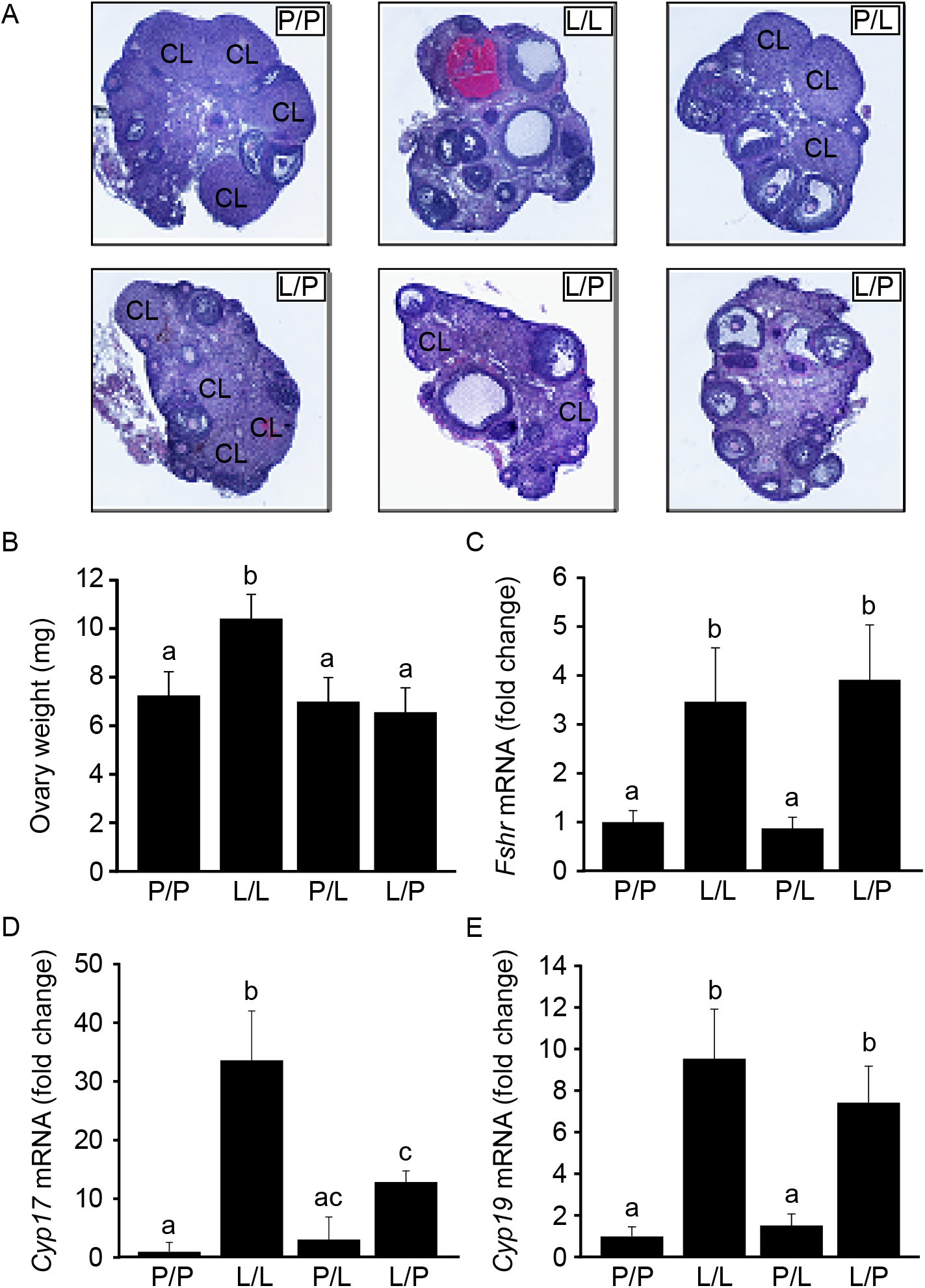
Co-housing letrozole mice with placebo mice improved the ovarian phenotype. The co-housing study included 3 groups of female mice housed 2 per cage: co-housed placebo mice (P/P), co-housed letrozole mice (L/L), and co-housed placebo with letrozole mice (P/L and L/P, respectively). Letrozole treatment (L/L) resulted in a lack of corpora lutea, cyst-like follicles and hemorrhagic cysts in the ovaries compared to ovaries from placebo mice (P/P) (A). Unlike L/L mice, L/P mice lacked polycystic ovaries and their ovaries contained corpora lutea (CL) which is evidence of ovulation (A). Letrozole (L/L) also resulted in increased ovarian weight and increased mRNA expression of several ovarian genes important in ovarian follicular development and steroidogenesis (B-E). Ovarian weight was lower in L/P mice compared to L/L mice (B). Follicle-stimulating hormone receptor *(Fshr)* and aromatase *(Cyp19)* mRNA levels were similar between L/L and L/P mice while cytochrome P450 17A1 *(Cyp17)* was decreased in L/P mice compared to L/L mice.

### Gut microbial richness did not correlate with an improved PCOS phenotype

The overall composition of the gut microbiome from samples collected before placebo and letrozole treatment (time 0) was compared amongst the four co-housing groups (P/P, L/L, P/L and L/P). No significant difference in alpha or beta diversity was observed among groups, indicating that the gut microbiomes were similar prior to treatment (Supplemental Fig. 1 A-B). Linear regression was used to examine the relationship between alpha diversity of the gut microbiome (Faith’s PD) and time. There was a strong positive relationship between alpha diversity and time in P/P mice (r = 0.23) but not L/L mice (r = 0.05) (Fig. 4A-B). To account for the repeated measures in this longitudinal study, we also used a linear mixed-effect model to examine the association between microbial diversity and time. This analysis confirmed that there was a significant effect of time on alpha diversity in P/P mice (p = 0.003) but not L/L mice (p = 0. 2) (Fig. 4A-B). These results agree with previous studies that reported lower alpha diversity of the gut microbiome in women with PCOS and rodent PCOS models compared to controls (24, 25, 27, 28). We then investigated whether changes in alpha diversity correlated with improved metabolic and reproductive phenotypes in L/P mice. In contrast to P/P mice, we did not observe a significant effect of time on alpha diversity using linear regression or the linear mixed-effect model in P/L (r = 0.009; p = 1) or L/P (r = 0.08; p = 0.71) mice (Fig. 4C-D). These results indicate that the physiological differences between L/L and L/P mice were probably not due to changes in alpha diversity *per se* but may reflect changes in specific gut microbes.

**Fig 4.**
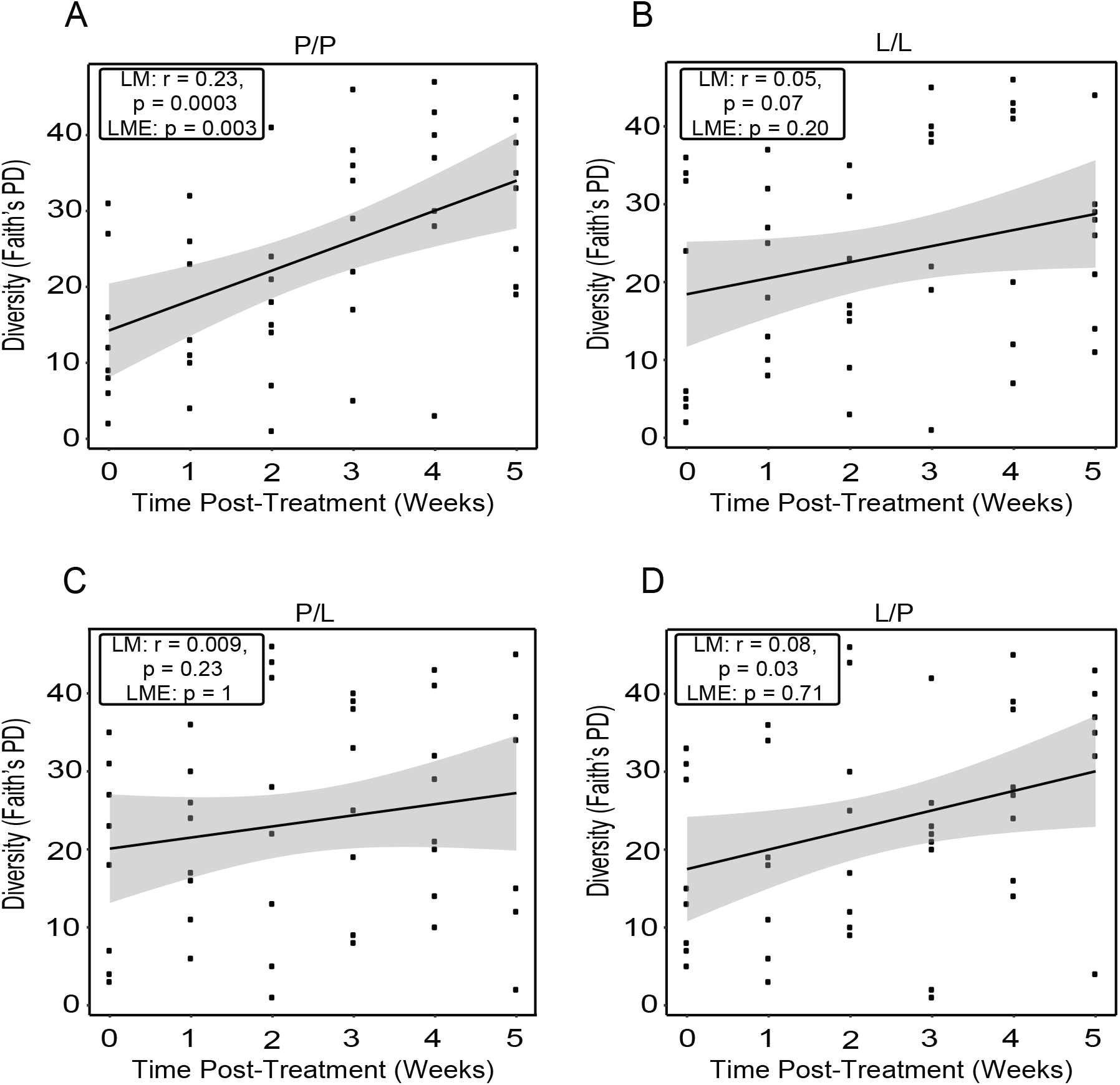
Co-housing letrozole mice with placebo mice did not restore alpha diversity of the gut microbiome. Alpha diversity as approximated by Faith’s phylogenetic diversity (PD) ranked estimate was graphed over time for co-housed placebo mice (P/P), co-housed letrozole mice (L/L), and co-housed placebo with letrozole mice (P/L and L/P, respectively). Results of linear regression model (LM) and P value are in the box inset, while the gray shaded area indicates the 95% confidence interval for the line of best fit. P values for the linear mixed effects model (LME) were obtained by the likelihood ratio test of the full model with the effect in question (time) against the model without the effect in question and are in the box inset.

### Composition of the gut microbiome was altered by co-housing

In addition to investigating changes in alpha diversity, we used weighted UniFrac distances to compare the similarity of gut microbial composition (beta diversity) amongst the different groups. While visualization of the UniFrac distances via Principal Coordinate Analysis (PCoA) did not result in distinct clustering, a PERMANOVA test (ADONIS) detected a significant effect of co-housing treatment on the microbial community structure (p=0.001) (Fig. 5A). This trend was also observed using unweighted UniFrac (data not shown). Canonical Analysis of Principal Coordinates (CAP) analysis was then used to analyze the microbial composition in response to an *a priori* defined experimental variable (co-housing treatment). PERMANOVA demonstrated a strong relationship between co-housing treatment and the overall composition of the gut microbiome (p=0.001) (Fig. 5B), suggesting that co-housing resulted in a distinct gut microbial community in L/P mice compared to L/L mice. To understand when the gut microbiome diverged, we then compared the fecal samples from the different co-housing treatment groups at each time point (Fig. 5C-G). By week 2, we observed a significant separation of the bacterial communities in the co-housing treatment groups (p = 0.004) (Fig. 5D). This separation occurred at the same time that we observed a protective effect of co-housing on weight gain in L/P mice (Fig. 1B), suggesting that changes in the abundance of specific bacteria may result in protection from developing the PCOS phenotype. It is notable that the CAP analysis indicated that co-housing also resulted in changes in the gut bacterial community of the P/L mice compared to P/P mice. However, since these changes were not sufficient to alter the metabolic and reproductive phenotypes of the host, these results suggest that the healthy gut microbiome was resistant to any pathological influence from the feces of the letrozole mice.

**Fig 5.**
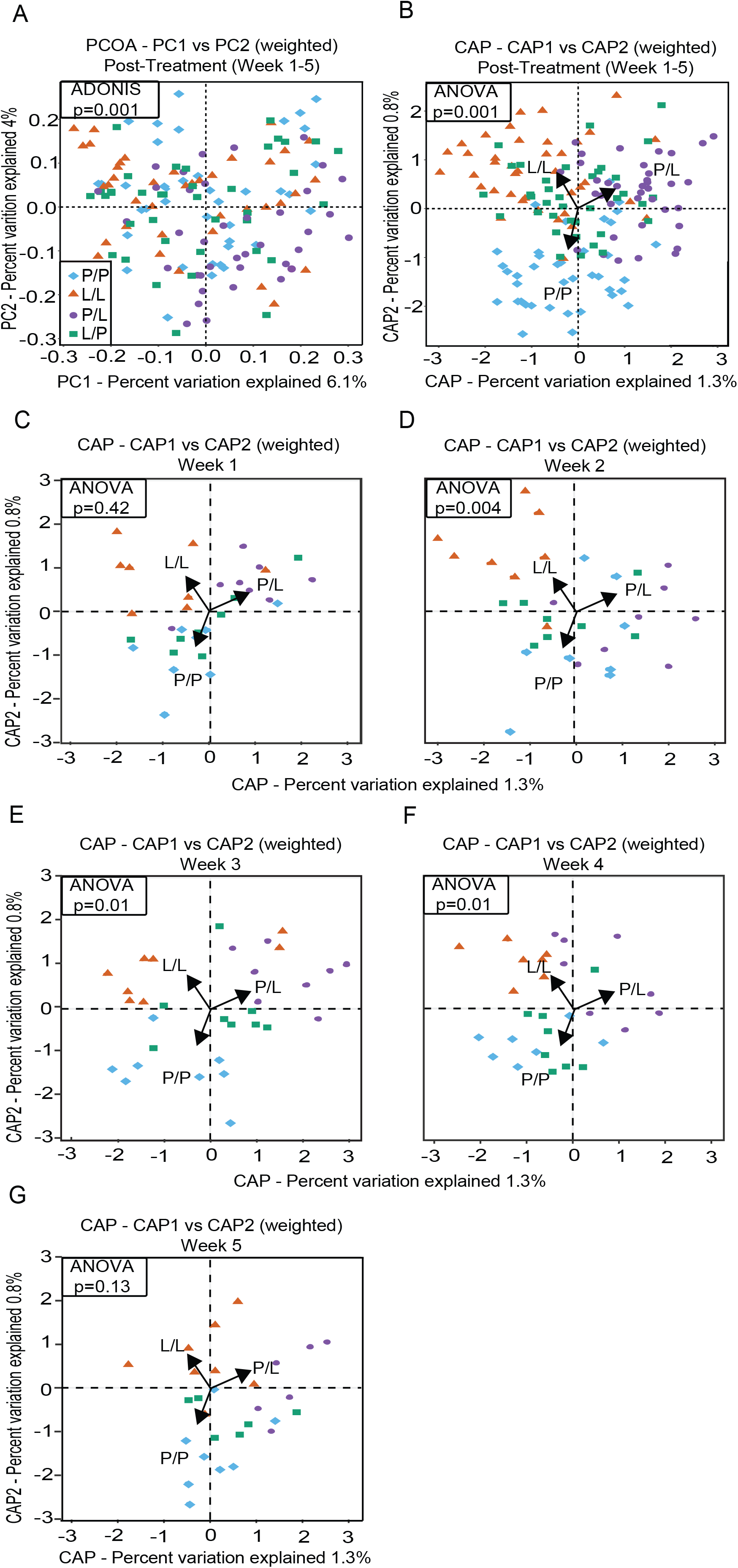
Co-housing letrozole mice with placebo mice influenced the overall composition of the gut bacterial community over time. Unconstrained principal coordinates analysis (PCoA) of weighted UniFrac distances demonstrated changes in the microbial composition (beta diversity) amongst samples collected post-treatment (A). Permutational analysis of variance of the weighted UniFrac distances indicated that co-housing had a strong influence on the gut microbial community (p=0.001). Constrained analysis of principal coordinates (CAP) of weighted UniFrac distances further illustrated the relationship between beta diversity and post-treatment with a significant effect of constraining the data based on co-housing treatment group (p=0.001) (B). Samples from the different co-housing groups were then compared at each time point (C-G). Permutational analysis of variance of the weighted UniFrac distances was done for each time point.

### Differentially abundant genera are associated with co-housed letrozole mice

Differential abundance of gut bacteria between placebo- and letrozole-treated mice was determined using DESeq2. This approach used a negative binomial regression for modeling count variables and is commonly used for overdispersed data, which is typical of microbiome data (31). DESeq2 identified five bacterial genera that were of higher relative abundance and four bacterial genera that were of lower relative abundance in placebo compared to letrozole mice (Fig. 6A). The gram-positive bacteria included *Coprobacillus*, *Candidatus Arthromitus*, *Roseburia*, *Dorea, Lactobacillus, Adlercreutzia,* and *Akkermansia* while the gram-negative bacteria included *Christensenella* and *Turicibacter*.

**Fig 6.**
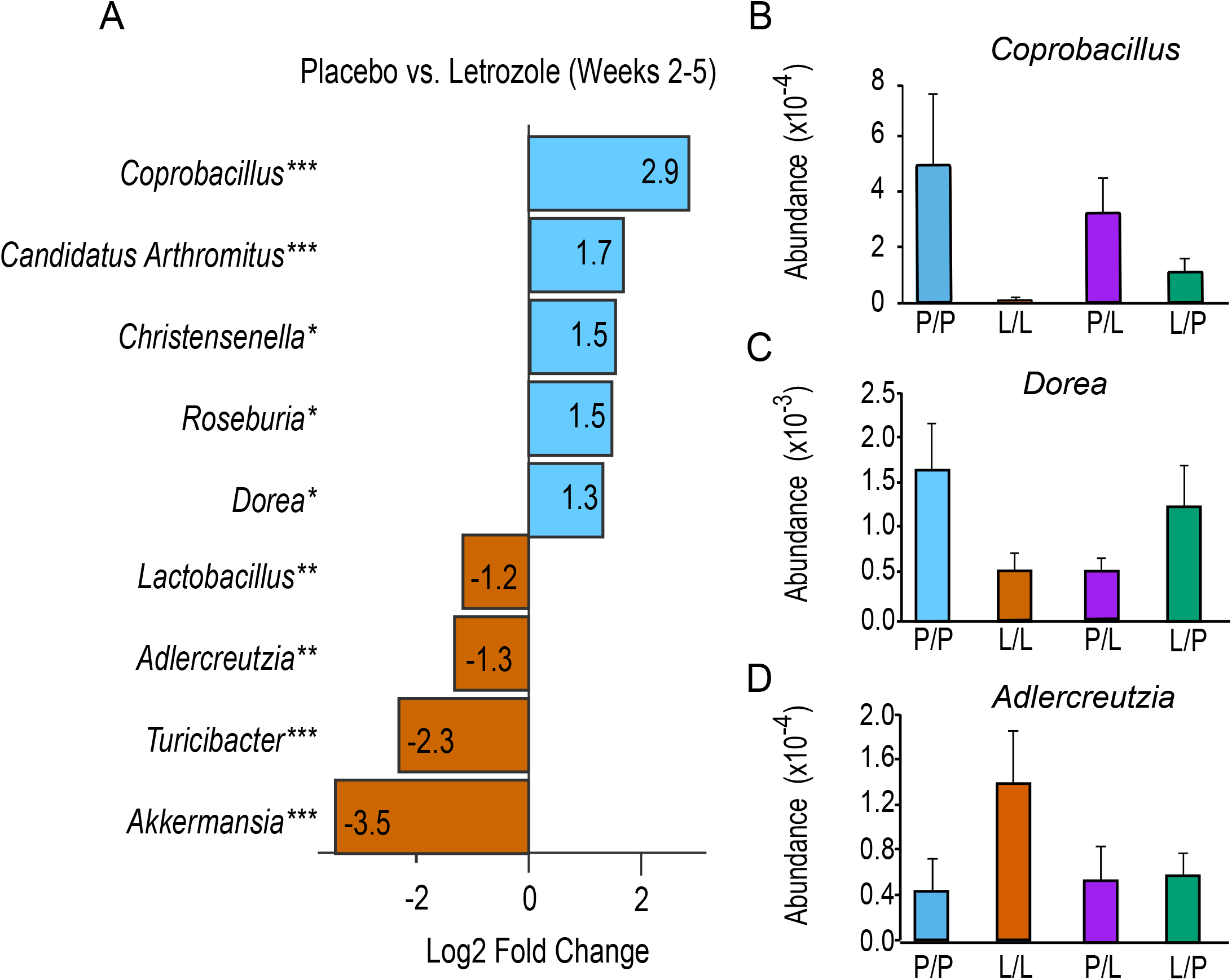
Specific bacterial genera were associated with improvement of the PCOS phenotype during co-housing. The co-housing study included 3 groups of female mice housed 2 per cage: co-housed placebo mice (P/P), co-housed letrozole mice (L/L), and co-housed placebo with letrozole mice (P/L and L/P, respectively). Results from the DESeq2 differential abundance analysis were expressed as log2 fold change for the comparison of L/L mice and P/P mice (A). Positive log2 fold changes represent bacterial genera increased in P/P mice relative to L/L mice while negative changes represent bacterial general increased in L/L mice relative to P/P mice. * p<0.05, ** p<0.01, *** p<0.001. Of the bacteria identified with the DESeq2 analysis, three bacterial genera had changes in relative abundance associated with co-housing letrozole mice with placebo mice (B-D).

From the DESeq2 analysis, we then identified three bacterial genera that had shifts in relative abundance corresponding with protection from developing the PCOS phenotype (Fig. 6B-D). For instance, the relative abundance of *Coprobacillus* was higher in both P/P and L/P mice compared to L/L mice. Interestingly, all three bacteria have been linked with host metabolism. *Coprobacillus* was reported to be enriched in healthy individuals compared to obese individuals and was proposed as a novel probiotic due to its association with a healthy gut microbiome (34, 35). *Dorea* was found to be more abundant in lean than obese individuals (36)and *Dorea longicatena* was included as a component of the RePOOPulate stool substitute being developed as a treatment for *Clostridium difficile* (37). Moreover, a higher abundance of *Adlercreutzia* was previously correlated with obesity (38). *Adlercreutzia* is also known to convert soy isoflavonoids to non-steroidal estrogens (equol) and was overrepresented in the guts of individuals who eat a soy diet (39).

In summary, our study demonstrated that co-housing letrozole mice with placebo mice resulted in protection from developing PCOS metabolic and reproductive phenotypes. In particular, exposure to a healthy gut microbiome resulted in letrozole-treated mice with body weight, FBG and insulin levels, and insulin resistance similar to placebo mice (Fig. 1). Surprisingly, co-housing letrozole-treated mice with healthy mice also resulted in letrozole-treated mice with testosterone and LH levels, estrous cycling and ovarian morphology similar to placebo mice (Fig. 2–3). This improvement in the PCOS phenotype correlated with changes in the relative abundance of specific gut bacteria such as *Coprobacillus*, *Dorea* and *Adlercreutzia*, indicating that manipulation of the gut microbiome towards a dysbiotic (letrozole treatment) or healthy state (co-housing) may influence the degree of pathology. Additional studies should characterize the effects of co-housing on the composition (strain level) and function of the gut microbiome in the letrozole-induced PCOS mouse model using metagenomics and metabolomics. Since these results imply that manipulating the composition of the gut microbiome may be a potential treatment option for women with PCOS, future studies should also investigate whether supplementation with prebiotics or novel probiotics such as *Coprobacillus* and *Dorea* can protect against development of PCOS.

## Materials and Methods

### PCOS Mouse Model

C57BL/6NHsd female mice from Envigo were housed in a vivarium under specific pathogen-free conditions with an automatic *12h:12h light/dark* cycle *(light* period: 06.00-18.00) and *ad libitum* access to water and food (Teklad S-2335 Mouse Breeder Irradiated Diet, Envigo). To establish the pubertal PCOS model, 4 week-old female mice were implanted subcutaneously with a placebo or 3 mg letrozole pellet (3 mm diameter; 50 μg/day; Innovative Research of America) for 5 weeks. For the co-housing paradigm, mice were housed two per cage: two placebo mice, two letrozole mice or one placebo and one letrozole mouse. All animal procedures in this study were approved by the University of California, San Diego Institutional Animal Care and Use Committee (protocol number S14011).

### Analysis of Reproductive Phenotype

Mice were weighed weekly. Estrous cycle stage was determined from the predominant cell type in vaginal epithelial smears obtained during weeks 4-5 of treatment as previously described (29). At the end of the experiment, ovaries were dissected, weighed, fixed in 4% paraformaldehyde, paraffin-embedded, sectioned at 10 μm and stained with hematoxylin and eosin (Zyagen). Serum testosterone and LH levels were measured using a mouse ELISA (range 10-800 ng/dL) and a radioimmunoassay (range 0.04-75 ng/mL), by the University of Virginia Center for Research in Reproduction Ligand Assay and Analysis Core Facility.

### Analysis of Metabolic Phenotype

After 5 weeks of treatment, mice were fasted for 5 hours and blood from the tail vein was collected to measure fasting insulin levels. Blood glucose was measured using a handheld glucometer (One Touch UltraMini, LifeScan, Inc) and an intraperitoneal (IP) insulin tolerance test (ITT) was performed. Tail vein blood glucose was measured just before (time 0) an IP injection of insulin (0.75 U/kg in sterile saline; Humulin R U-100, Eli Lilly) was given and at 15, 30, 45, 60, 90, and 120 minutes post injection. At the end of the experiment, the mice were anesthetized with isoflurane, blood was collected from the posterior vena cava, and parametrial fat pads were dissected and weighed. Serum insulin was measured using a mouse ELISA (ALPO) by the University of California, Davis Mouse Metabolic Phenotyping Center. Differences among groups were analyzed by one-way ANOVA or two-way repeated measures ANOVA followed by post-hoc comparisons of individual time points.

### Quantitative Real-Time Polymerase Chain Reaction of Ovarian Genes

Total RNA was isolated from ovaries using a RNeasy Mini Plus kit (Qiagen), which also removes genomic DNA. Complementary DNA (cDNA) was made by reverse transcription of total RNA using an iScript cDNA synthesis kit (Bio-Rad Laboratories). cDNA products were detected using SYBR Green Supermix (Bio-Rad Laboratories) on a Bio-Rad CFX Connect quantitative real-time polymerase chain reaction system (Bio-Rad Laboratories) using previously described primers (29). Data were analyzed by the 2^2ΔΔCt^ method (40) by normalizing the gene of interest to GAPDH. Data were represented as mean fold change compared with placebo ± the standard error of the mean.

### Fecal Sample Collection, DNA Isolation and 16S rRNA Gene Sequencing

Fecal samples were collected prior to treatment and once per week for 5 weeks. Fecal samples were frozen immediately after collection and stored at -80°C. Bacterial DNA was extracted from the samples using the DNeasy PowerSoil Kit (Qiagen) and stored at -80°C. The V4 hypervariable region of the 16S rRNA gene was PCR amplified with primers 515F and 806R (41). The reverse primers contained unique 12 base pair Golay barcodes that were incorporated into the PCR amplicons (41). Amplicon sequence libraries were prepared at The Scripps Research Institute Next Generation Sequencing Core Facility where the libraries were sequenced on an Illumina MiSeq as previously described (27).

### 16S rRNA Gene Sequence Analysis

Raw sequences were imported into QIIME 2 (version 2018.4) using the q2-tools-import script and sequences were demultiplexed using the q2-demux emp-single script. This resulted in 7.3 million sequences with an average of 36,000 sequences per sample. The 16S rRNA sequences generated in this study were deposited into the European Nucleotide Archive (Study Accession Number PRJEB29583). DADA2 software was used to obtain a set of Observed Sequence Variants (SVs) (42). Based on the quality scores, the forward reads were truncated at position 240 using the q2-dada2-denoise script. Taxonomy was assigned using a pre-trained Naïve Bayes classifier (Greengenes 13_8 99% operational taxonomic units) and the q2-feature-classifier plugin (43). In total, 318 SVs were identified from 186 fecal pellets. The resulting SVs were then aligned using MAFFT (44) and a phylogenetic tree was built using FastTree (45). Taxonomic distributions of the samples were calculated using the q2-taxa-barplot script. Alpha and beta diversity metrics were computed using the q2-diversity core-metrics script at a rarefied sampling depth of 1250. Rarefaction resulted in the removal of one sample that had ~400 sequences. The alpha diversity metric, Faith’s Phylogenetic Diversity (PD), was used to measure phylogenetic biodiversity by calculating the total branch lengths on a phylogenetic tree of all members in a community (46). UniFrac was used to compare the similarity (beta diversity) among the microbial communities by calculating the shared PD between pairs of microbial communities (47, 48).

### Statistical Analysis

Statistical calculations were performed in the R Studio statistical package (version 0.99.893) with the phyloseq (version 1.19.1) (49) and vegan package (version 2.5.2). Alpha diversity data was tested for normality via the Shapiro-Wilk test. Variables that were not normally distributed were ranked. Changes in alpha diversity over time were analyzed using simple linear regression and Pearson’s rank correlation on ranked diversity measures. Linear mixed effects analysis of the relationship between alpha diversity and time was done with the lme4 R package (version 1.1.18.1). P-values were obtained by likelihood ratio tests of the full model with the effect in question against the model without the effect in question. Principal Coordinate Analysis (PCoA) and CAP plots (50) were constructed using the phyloseq R package. PCoA plots were used to represent the similarity of post-treatment (Weeks 1-5) fecal microbiome samples based on multiple variables in the data set, while CAP was used to visualize the relationship of the fecal microbiome with specific parameters. Permutational multivariate analysis of variance (PERMANOVA) used post-treatment weighted UniFrac distance measures to assess bacterial community compositional differences and its relationship to co-housing treatment group (999 permutations “vegan” package). DESeq2 (51) (version 1.14.1) in the microbiomeSeq package (version 0.1 http://www.github.com/umerijaz/microbiomeSeq) was used to identify bacterial genera that were differentially abundant between placebo and letrozole-treated mice.

## List of Abbreviations

FBG: fasting blood glucose
ITT: insulin tolerance test
IP: intraperitoneal
SV: sequence variant
PERMANOVA: permutational multivariate analysis of variance
PD: phylogenetic diversity
LH: luteinizing hormone
CAP: canonical analysis of principal coordinates
PCoA: principal coordinates analysis
PCOS: polycystic ovary syndrome
QIIME: Quantitative Insights Into Microbial Ecology

## Acknowledgements

We thank members of the Kelley and Thackray labs for insightful comments and suggestions. Hormone levels were measured by the University of Virginia Center for Research in Reproduction Ligand Assay and Analysis Core Facility (P50 HD28934) and the University of California, Davis Mouse Metabolic Phenotyping Core (U24 DK092993). This work was funded by the National Institute of Child Health and Human Development through a cooperative agreement as part of the National Centers for Translational Research in Reproduction and Infertility (P50 HD012303). P.J.T. was also funded by the UCSD Microbial Sciences Research Initiative Fellowship and the SDSU ARCS Foundation. L.S. and A.C. were funded by a Doris A. Howell Foundation Research Scholarship for Women’s Health.

## Authors’ contributions

V.G.T. and S.T.K. conceived and designed the study; P.J.T., B.S.H., P.A., L.S., and A.C. collected samples, performed reproductive and metabolic assessments, quantitative RT-PCR; P.J.T. performed DNA extractions and PCR amplifications; P.J.T., V.G.T. and S.T.K. analyzed the data; P.J.T., S.T.K. and V.G.T. wrote the manuscript.

## References

1. Fauser BC, et al. (2012) Consensus on women’s health aspects of polycystic ovary syndrome (PCOS): the Amsterdam ESHRE/ASRM-Sponsored 3rd PCOS Consensus Workshop Group. Fertil Steril 97(1):28–38 e25.

2. Azziz R, et al. (2016) Polycystic ovary syndrome. Nat Rev Dis Primers 2:16057.

3. Churchill SJ, Wang ET, & Pisarska MD (2015) Metabolic consequences of polycystic ovary syndrome. Minerva Ginecol 67(6):545–555.

4. Goodman NF, et al. (2015) American Association of Clinical Endocrinologists, American College of Endocrinology, and Androgen Excess and Pcos Society Disease State Clinical Review: Guide to the Best Practices in the Evaluation and Treatment of Polycystic Ovary Syndrome -Part 2. Endocrine Practice 21(12):1415–1426.

5. Barber TM, Wass JAH, McCarthy MI, & Franks S (2007) Metabolic characteristics of women with polycystic ovaries and oligo-amenorrhoea but normal androgen levels: implications for the management of polycystic ovary syndrome. Clinical endocrinology 66(4):513–517.

6. Moghetti P, et al. (2013) Divergences in Insulin Resistance Between the Different Phenotypes of the Polycystic Ovary Syndrome. The Journal of Clinical Endocrinology & Metabolism 98(4):E628–E637.

7. Barber TM, Wass JA, McCarthy MI, & Franks S (2007) Metabolic characteristics of women with polycystic ovaries and oligo-amenorrhoea but normal androgen levels: implications for the management of polycystic ovary syndrome. Clin Endocrinol (Oxf) 66(4):513–517.

8. Legro RS, Driscoll D, Strauss JF, Fox J, & Dunaif A (1998) Evidence for a genetic basis for hyperandrogenemia in polycystic ovary syndrome. Proc. Natl. Acad. Sci. 14956–14960.

9. Vink JM, Sadrzadeh S, Lambalk CB, & Boomsma DI (2006) Heritability of polycystic ovary syndrome in a Dutch twin-family study. J Clin Endocrinol Metab 91(6):2100–2104.

10. Abbott DH & Bacha F (2013) Ontogeny of polycystic ovary syndrome and insulin resistance in utero and early childhood. Fertil Steril 100(1):2–11.

11. Clemente JC, Ursell LK, Parfrey LW, & Knight R (2012) The impact of the gut microbiota on human health: an integrative view. Cell 148(6):1258–1270.

12. Walker AW & Lawley TD (2013) Therapeutic modulation of intestinal dysbiosis. (Translated from eng) Pharmacol Res 69(1):75–86 (in eng).

13. Baumler AJ & Sperandio V (2016) Interactions between the microbiota and pathogenic bacteria in the gut. Nature 535(7610):85–93.

14. Gensollen T, Iyer SS, Kasper DL, & Blumberg RS (2016) How colonization by microbiota in early life shapes the immune system. Science 352(6285):539–544.

15. Natividad JM & Verdu EF (2013) Modulation of intestinal barrier by intestinal microbiota: pathological and therapeutic implications. Pharmacol Res 69(1):42–51.

16. den Besten G, et al. (2013) The role of short-chain fatty acids in the interplay between diet, gut microbiota, and host energy metabolism. J Lipid Res 54(9):2325–2340.

17. Ridlon JM, Kang DJ, Hylemon PB, & Bajaj JS (2014) Bile acids and the gut microbiome. Curr Opin Gastroenterol 30(3):332–338.

18. Ley RE, Turnbaugh PJ, Klein S, & Gordon JI (2006) Microbial ecology: human gut microbes associated with obesity. Nature 444(7122):1022–1023.

19. Qin J, et al. (2012) A metagenome-wide association study of gut microbiota in type 2 diabetes. Nature 490(7418):55–60.

20. Turnbaugh PJ, et al. (2009) A core gut microbiome in obese and lean twins. Nature 457(7228):480–484.

21. Turnbaugh PJ, et al. (2006) An obesity-associated gut microbiome with increased capacity for energy harvest. Nature 444(7122):1027–1031.

22. Ridaura VK, et al. (2013) Gut microbiota from twins discordant for obesity modulate metabolism in mice. Science 341(6150):1241214.

23. Liu R, et al. (2017) Dysbiosis of Gut Microbiota Associated with Clinical Parameters in Polycystic Ovary Syndrome. Frontiers in Microbiology 8.

24. Lindheim L, et al. (2017) Alterations in Gut Microbiome Composition and Barrier Function Are Associated with Reproductive and Metabolic Defects in Women with Polycystic Ovary Syndrome (PCOS): A Pilot Study. PLoS One 12(1):e0168390.

25. Torres PJ, et al. (2018) Gut Microbial Diversity in Women with Polycystic Ovary Syndrome Correlates with Hyperandrogenism. J Clin Endocrinol Metab.

26. Insenser M, et al. (2018) Gut Microbiota and the Polycystic Ovary Syndrome: Influence of Sex, Sex Hormones, and Obesity. J Clin Endocrinol Metab 103(7):2552–2562.

27. Kelley ST, Skarra DV, Rivera AJ, & Thackray VG (2016) The Gut Microbiome Is Altered in a Letrozole-Induced Mouse Model of Polycystic Ovary Syndrome. PLoS One 11(1):e0146509.

28. Guo YJ, et al. (2016) Association between Polycystic Ovary Syndrome and Gut Microbiota. Plos One 11(4).

29. Kauffman AS, et al. (2015) A novel letrozole model recapitulates both the reproductive and metabolic phenotypes of Polycystic Ovary Syndrome in female mice. Biol Reprod 93(3):69.

30. Skarra DV, Hernandez-Carretero A, Rivera AJ, Anvar AR, & Thackray VG (2017) Hyperandrogenemia Induced by Letrozole Treatment of Pubertal Female Mice Results in Hyperinsulinemia Prior to Weight Gain and Insulin Resistance. Endocrinology 158(9):2988–3003.

31. Maier L, et al. (2018) Extensive impact of non-antibiotic drugs on human gut bacteria. Nature 555(7698):623–628.

32. Kumar A, Magoffin D, Munir I, & Azziz R (2009) Effect of insulin and testosterone on androgen production and transcription of SULT2A1 in the NCI-H295R adrenocortical cell line. Fertil Steril 92(2):793–797.

33. Zhang G & Veldhuis JD (2004) Insulin drives transcriptional activity of the CYP17 gene in primary cultures of swine theca cells. Biol Reprod 70(6):1600–1605.

34. Hou YP, et al. (2017) Human Gut Microbiota Associated with Obesity in Chinese Children and Adolescents. Biomed Res Int 2017:7585989.

35. Tap J, et al. (2009) Towards the human intestinal microbiota phylogenetic core. Environ Microbiol 11(10):2574–2584.

36. Aguirre M, Bussolo de Souza C, & Venema K (2016) The Gut Microbiota from Lean and Obese Subjects Contribute Differently to the Fermentation of Arabinogalactan and Inulin. PLoS One 11(7):e0159236.

37. Petrof EO, et al. (2013) Stool substitute transplant therapy for the eradication of Clostridium difficile infection: ‘RePOOPulating’ the gut. Microbiome 1(1):3.

38. Yun Y, et al. (2017) Comparative analysis of gut microbiota associated with body mass index in a large Korean cohort. BMC Microbiol 17(1):151.

39. Rafii F (2015) The role of colonic bacteria in the metabolism of the natural isoflavone daidzin to equol. Metabolites 5(1):56–73.

40. Livak KJ & Schmittgen TD (2001) Analysis of relative gene expression data using real-time quantitative PCR and the 2(-Delta Delta C(T)) Method. Methods 25(4):402–408.

41. Caporaso JG, et al. (2012) Ultra-high-throughput microbial community analysis on the Illumina HiSeq and MiSeq platforms. ISME J 6(8):1621–1624.

42. Callahan BJ, et al. (2016) DADA2: High-resolution sample inference from Illumina amplicon data. Nat Methods 13(7):581–583.

43. Bokulich NA, et al. (2018) Optimizing taxonomic classification of marker-gene amplicon sequences with QIIME 2’s q2-feature-classifier plugin. Microbiome 6(1):90.

44. Katoh K & Standley DM (2013) MAFFT multiple sequence alignment software version 7: improvements in performance and usability. Mol Biol Evol 30(4):772–780.

45. Price MN, Dehal PS, & Arkin AP (2009) FastTree: computing large minimum evolution trees with profiles instead of a distance matrix. Mol Biol Evol 26(7):1641–1650.

46. Faith DP (1992) Conservation evaluation and phylogenetic diversity. Biological Conservation 61(1):1–10.

47. Lozupone C & Knight R (2005) UniFrac: a new phylogenetic method for comparing microbial communities. Appl Environ Microbiol 71(12):8228–8235.

48. Lozupone C, Lladser ME, Knights D, Stombaugh J, & Knight R (2011) UniFrac: an effective distance metric for microbial community comparison. ISME J 5(2):169–172.

49. McMurdie PJ & Holmes S (2012) Phyloseq: a bioconductor package for handling and analysis of high-throughput phylogenetic sequence data. Pac Symp Biocomput:235–246.

50. Anderson MJ & Willis TJ (2003) Canonical analysis of principal coordinates: A useful method of constrained ordination for ecology. Ecology 84(2):511–525.

51. Love MI, Huber W, & Anders S (2014) Moderated estimation of fold change and dispersion for RNA-seq data with DESeq2. Genome Biol 15(12):550.

